# Mechanisms of high-intensity sound exposure on inhibiting hippocampal long-term potentiation: role of brain-derived neurotrophic factor

**DOI:** 10.1101/850214

**Authors:** Júnia L. de Deus, Mateus R. Amorim, Aline B. Ribeiro, Procópio C. G. Barcellos-Filho, César C. Ceballos, Luiz Guilherme S. Branco, Alexandra O.S. Cunha, Ricardo M. Leão

## Abstract

Exposure to humans and experimental animals to loud noises produce cognitive and emotional disorders and recent studies have shown that hippocampal neuronal function is affected by auditory stimulation or deprivation. We have found previously that in the hippocampus of rats exposed to high-intensity sound (110 dB) for one-minute the Schaffer-CA1 long-term potentiation (LTP) is strongly inhibited. Here we investigated possible mechanisms involved in this effect. We found, using c-fos expression, that exposure to 110 dB sound-activated neurons in the CA1 and CA3 hippocampal region. Using electrophysiological recordings in hippocampal slices, we found that both GABAergic and glutamatergic neurotransmission were unaffected by high-intensity sound stimulation. However, hippocampal brain-derived neurotrophic factor (BDNF), which is involved in promoting hippocampal synaptic plasticity, presented decreased levels in sound-stimulated animals. Perfusion of slices with BDNF revert the inhibition of LTP after a single sound stimulus in comparison to sham-stimulated rats. Furthermore, the perfusion with LM 22A4, a TrkB receptor agonist also rescued LTP from sound-stimulated animals. Our results strongly suggest that the exposure to high-intensity sound inhibits the BDNF production in the hippocampus, which could be a possible mechanism of the inhibition of LTP by high-intensity sound exposure.

## Introduction

Loud sounds can damage cochlear hair cells and is currently the leading cause of permanent hearing loss and tinnitus (Le et al., 2017). Even prolonged exposure to non-traumatic noise has adverse effects on the auditory processing (Eggermont et al., 2017) and exposure to moderate to high-intensity sound triggers behavior responses resulting in the avoidance of the sound source (Manohar et al., 2017), a protective measure to avoid damage to auditory hair cells. Indeed, loud sounds are stressors that lead to the activation of the HPA axis, leading to increased secretion of corticosterone (Helfferich and Palkovits, 2003; Burrow et al., 2005). Thus, non-auditory areas are affected by the level of auditory input and adjust the animal’s behavior to the acoustic environment.

Additionally, exposure to loud and even moderate sounds can produce several systemic and cognitive deficits in humans (Lercher et al., 2003; Stansfeld et al., 2005; Basner et al., 2014). The so-called “sonic attack” to the American and Canadian diplomatic personnel in Havana, Cuba, which produced several emotional and cognitive symptoms after the exposure to a high-pitched loud sound (Hofner et al., 2018; Swanson et al., 2018), is a dramatic example of the potential harm that loud sound exposure can accomplish beyond the auditory system.

The hippocampus receives sensory information from several modalities, which are fundamental for its role in spatial navigation, learning and memory formation (Save et al., 2000; Jeffery, 2007; Ravassard et al., 2013). It is connected to the auditory system (Kraus and Canlon, 2012; Zhang et al., 2018) and sound stimulation triggers excitatory and inhibitory neurotransmission in hippocampal neurons *in vivo* (Wang et al., 2017). Also, the hippocampus is related to the formation of auditory memories (Squire et al., 2001), it uses auditory information for the formation of spatial memory (Tamura et al., 1990) and hippocampal place cells can be activated by auditory dimension cues (Aronov et al., 2017). On the other hand, prolonged exposure to loud or even moderate-intensity sounds can affect the hippocampus negatively. In this case, prolonged exposure to moderate-intensity sound increases oxidative damage and tau phosphorylation in the hippocampus and impairs spatial memory in mice (Cheng et al., 2011,2016), whereas high-intensity sound exposure affects hippocampal place cells activity (Goble et al., 2009). Additionally, exposure to moderate to loud sounds can lead to impairment of spatial and associative memory, which seems to be associated to an oxidative status imbalance (Uran et al., 2010, 2012; Manikandan et al., 2016). On the other hand, 40-minutes exposure to mild (80dB) sounds, potentiated hippocampal long-term potentiation (LTP) and brain-derived neurotrophic factor (BDNF) expression (Matt et al., 2018). Finally, we have shown that one minute exposure to a high-intensity (110-120 dB) broadband sound inhibits LTP in the Schaffer-CA1 synapses in the hippocampus of rats for 24 hours after exposure (de Deus et al., 2017), an effect not related to corticosterone secretion, and not observed after moderate noise (80 dB) exposure.

Hippocampal LTP is a long-lasting enhancement of synaptic efficacy associated with hippocampal learning and memory (Bliss and Lomo, 1973; Bliss and Collingridge, 1993). In the Schaffer-CA1 synapse, the LTP is dependent on post-synaptic calcium influx via glutamatergic NMDA receptors (for review see Malenka and Bear, 2004; Nicoll, 2017). LTP is modulated positively or negatively by several signaling molecules, such as BDNF, a neurotrophic factor which acts on TrkB receptors (Minichielo, 2009). Activation of TrkB receptors by BDNF promotes LTP in the Schaffer-CA1 synapse (Minichielo, 2009; Edelman et al., 2014; Lin et al., 2018). Heterozygous BDNF (+/−) knockout mice have a significant deficit in hippocampal LTP (Korte et al., 1995), which can be rescued by exogenous BDNF (Paterson et al., 1996). The secretion of BNDF is also promoted by stimulation protocols that develop LTP in contrast to stimulation patterns that do not lead to LTP (Aicardi et al., 2004). In accordance to the role of BDNF in facilitating the induction of LTP, BDNF can revert LTP in situations where the LTP is inhibited by exogenous conditions, like after prolonged periods of chronic intermittent hypoxia (Xie et al., 2010).

Here we investigated the mechanisms of LTP inhibition by a single episode of high-intensity sound stimulation (de Deus et al., 2017). We demonstrated that both glutamatergic and GABAergic transmission is unaffected by a single episode of high intense sound stimulation, but the levels of BDNF in the hippocampus are reduced, and LTP could be rescued by exogenous BDNF application.

## Materials and Methods

All experimental procedures involving animals were elaborated according to the rules of research in the National Council for Control of Animal Experimentation and approved by the Committee on Ethics in Animal Use of the Ribeirão Preto Medical School of the University of São Paulo, (protocol # 006/2-2015).

### Animals

Experiments were performed on male Wistar rats 60-70 days old obtained from the Central Animal Facility of the Ribeirão Preto Campus of the University of São Paulo. Rats were kept in Plexiglas cages (2-3 animals per cage) at the Medical School of Ribeirão Preto, with food and water available *ad libitum* and a 12-hr dark/light cycle (lights on at 7:00 a.m.) and controlled temperature (22°C). We divided the animals in two experimental groups: sham-stimulated (rats placed in the sound stimulus box without sound stimulus) and rats submitted to a sound stimulus of 110 dB (high-intensity sound).

### Sound stimulation

Rats were placed in an acrylic arena (height: 32 cm, diameter: 30 cm), located inside an acoustically isolated chamber (45 × 45 × 40 cm), with 2 loudspeakers placed at the top of the arena, in accordance with protocols described previously by de Deus et al., (2017). After oneminute acclimatization, they were submitted to a one-minute duration stimulus of 110 dB, consisting of a digitally modified recording of a doorbell, spanning frequencies from 3 to 15 kHz (Romcy-Pereira and Garcia-Cairasco, 2003). After stimulation, animals were kept in the arena for one more minute and returned to their boxes where they remained for two hours until the preparation of the hippocampal slices. Sham-stimulated animals were placed in the same arena for 3 minutes without any sound stimulus. Ambient noise inside the acoustic chamber was 55 dB, and the sound intensity inside the arena was checked and calibrated regularly with a decibel meter (Extech 407730 - Sound Level Meter).

### c-Fos Immunofluorescence

After 90 minutes of the sound or sham stimulus, rats were anesthetized with isoflurane for transcardial perfusion with phosphate buffered saline (PBS, 0.1 M, pH 7.4) and paraformaldehyde at 4% dissolved in PBS. After perfusion, the brains were immediately removed and immersed in 30% sucrose solution in PBS until tissue saturation. Next, tissue blocks were frozen in isopentane for 30 s and stored at −80 °C until sectioned. Coronal sections of 40 μm were cut, in a cryostat, in four series in the CA1 region. The sections were washed rapidly with glycine (0.1M) to remove excess paraformaldehyde, permeabilized with 0.1% Triton X-100 for 10 minutes and washed with 0.1M PBS. Nonspecific binding was blocked with PBS containing 1% bovine serum albumin (BSA; Sigma, St. Louis, MO, USA) for 1 hour. Subsequently, sections were incubated for 40 hours at 4°C with a polyclonal primary anti-c-Fos antibody produced in rabbit (1: 5000; Santa Cruz Biotechnology, Dallas, TX, USA). The primary antibody was diluted in PBS containing 0.1% Triton X-100 and 1% BSA. Then, after washing with PBS (5X for 5 min), sections were incubated with biotinylated anti-goat antibody IgG produced in rabbit (1: 1000; Alexa 594, Vector Laboratories, Burlingame, CA, USA) for 1 hour. Finally, nuclear labeling was done with DAPI (4’, 6-diamidino-2-phenylindole; 1 μg/mL, Sigma-Aldrich) for 10 minutes at room temperature. Sections were then mounted on slides using Fluoromount G (Electron Microscopy Sciences, Hatfield, PA) as assembling medium, viewed and photographed under fluorescence microscopy DM 5500B (Leica Microsystems, Wetzlar, Germany). Negative control of the primary antibody was performed, where they were omitted, and the cuts were only incubated with the secondary antibodies. No labeling was observed on these controls.

### Preparation of Hippocampal Slices

After two hours of the sound or sham stimulus, the animals were anesthetized with isoflurane and decapitated. The brains were rapidly removed and placed in an ice-cold solution containing: 87 NaCl, 2.5 KCl, 25 NaHCO_3_, 1.25 NaH_2_PO_4_, 75 sucrose, 25 Glucose, 0.2 CaCl_2_, 7 MgCl_2_, bubbled with 95% O_2_ and 5% CO_2_. Brain hemispheres were separated, positioned side by side, fixed with cyanoacrylate glue (SuperBonder^®^) to a base and placed in the vibrating chamber of a vibratome (1000 plus, Vibratome, USA). Transversal slices containing the dorsal hippocampus (200 μm for whole-cell recordings and 400 μm for extracellular recordings) cut and the hippocampus was separated from the cortex using ophthalmic scissors and microtweezers. Slices were placed in artificial cerebrospinal fluid (aCSF) solution containing (mM): 125 NaCl, 2.5 KCl, 1.25 NaH_2_PO_4_, 26 NaHCO_3_, 10 Glucose, 2 CaCl_2_, 1 MgCl_2_ and left to rest for at least two hours before use (one hour at 34-35 °C and at least one hour in room temperature) continuously bubbled with a carbogenic mixture (95% O_2_ and 5% CO_2_).

### Whole-Cell Patch-Clamp Recordings

Whole-cell patch-clamp experiments were performed in slices placed in a chamber continuously perfused aCSF (1 ml/min) supplemented with picrotoxin (GABA_A_ receptor antagonist: 20 μM) except for GABAergic neurotransmission, and at a temperature of 34 °C controlled with inline heater (Scientifica). Pyramidal neurons from the CA1 region were visualized with an Olympus BX51WI microscope through a 40x water immersion objective and with infrared differential interference contrast (IR-DIC). Pyramidal cells were chosen according to the morphology (pyramidal shape) and position, in the middle of the pyramidal layer. Recording electrodes were filled with an internal pipette solution chosen for each set of experiments and fabricated from borosilicate capillaries (BF150-86-10, 15 Sutter Instruments) with the tip resistance between 4-5 MΩ. Electrophysiological recordings in whole-cell patchclamp were performed using a Heka EPC10 amplifier, with a sampling rate of 50 kHz and low pass filtered at 3 kHz (Bessel). Neurons that presented series resistance increased over 20% during experiments, as well as resting membrane potential higher than −60 mV, were discarded. Series resistance (<20 MΩ) was compensated at around 60%. Voltages were corrected off-line for a liquid junction potential for each internal solution calculated with Clampex software (Molecular Devices).

Glutamatergic EPSCs were evoked from the stimulation of the Schaffer pathway with a concentric bipolar microelectrode (FHC-Bowdoin, ME, USA) connected to an SD9 Grass voltage stimulator (Natus Medical Incorporated, Warwick, RI, USA). AMPA and NMDA-EPSCs recording were performed under the perfusion of the slices with picrotoxin (20 μM) using an internal solution composed in mM: 130 CsCl, 10 HEPES, 5 EGTA, 5 phosphocreatine, 4 Mg-ATP, 0.5 Na-GTP, 10 TEA, 5 QX 314, adjusted to pH 7.3 with CsOH and ≈290 mOsm/kgH_2_O. AMPA and NMDA currents were recorded at potentials from −70 to +80 mV with increments of +30 mV. The stimulus intensity was gradually increased until the amplitude of the synaptic current reached its maximum. We then stimulated Schaffer pathway with the minimum voltage necessary to evoke a maximum post-synaptic current for recording the currents. To obtain NMDA currents, we blocked AMPA/KA currents with DNQX (10 μM) for 10 minutes and to estimate AMPA currents, we subtracted currents before and after DNQX.

Miniature GABAergic IPSCs (mIPSCs) were recorded for 10 minutes at −70 mV with an internal solution consisting in (mM): 145 KCl, 10 HEPES, 0.5 EGTA, 10 phosphocreatine, 4 Mg-ATP, 0.3 Na-GTP, adjusted to pH 7.3 with KOH and ≈290 mOsm/kg H_2_O. Miniature IPSCs (mIPSC) were recorded in aCSF in the presence of TTX (0.5 μM) and DNQX (10 μM).

### BDNF quantification

BNDF was quantified using ELISA. In these experiments, 4 hours after sham or sound stimulus, rats were decapitated and the brains were carefully collected, frozen by submersion in dry ice-cold isopentane and kept under −80 °C. The dorsal hippocampus was sampled in a cryostat by a punch needle (1.5 mm inner diameter) from a 1200-μm thick slice (Figure 4A) and stored in plastic tubes at −80 °C until analysis. Samples were homogenized with buffer solution (137 mM NaCl, 20 mM Tris-HCl pH 7.6, 10% glycerol and sodium orthovanadate), supplemented with phosphatase protease inhibitor cocktail (Cell Signaling, Massachusetts, USA) and centrifuged at 13.000 rpm for 20 min at 4 °C. Tissue supernatant were used to estimate BDNF (detection limits 7.8-500 pg/ml) by ELISA (BDNF Emax Immuno Assay System, #G7611, Promega), according to the manufacturer’s instructions. Results from BDNF in hippocampal homogenates were normalized by protein concentrations, which were assessed by the Bradford assay (#5000205, Bio-Rad Laboratories, USA).

### Field potential recordings and LTP induction

Extracellular electrophysiological recordings were performed with a Multiclamp 700B amplifier (Molecular Devices, USA) connected to a Digidata 1440 A AC/DC interface (Molecular Devices, USA). Slices were placed in the recording chamber with continuous superfusion of aCSF (1 mL/min) bubbled with a carbogenic mixture and the temperature-controlled (32-34 °C) using an inline temperature heater (Warner Instruments, USA). To stimulate Schaffer/collateral fibers, we used a stainless steel bipolar concentric microelectrode (FHC-Bowdoin, Maine, USA) connected to a Master-9 voltage stimulator (A.M.P.I., Israel). Field excitatory post-synaptic potentials (fEPSPs) were recorded at CA1 *stratum radiatum* with borosilicate glass microelectrodes (G85150T, Warner Instruments, USA) filled with aCSF with tip resistances of 1-2 MΩ. For LTP induction, first, we performed an input-output curve, where the voltage was gradually increased by 10 V increments until population spikes were observed in the fEPSP. From the maximum stimulus, we choose the stimulus intensity that produced a fEPSP equivalent to approximately 50% of the maximum response. Subsequently, 50 fEPSPs were recorded at this intensity, for 25 minutes at 0.03 Hz. After baseline recording, LTP was induced on Schaffer-collateral fibers using 3 trains of high-frequency stimulation (HFS) at 100 Hz, 1-second duration (inter-train interval of 3 s) and after that, we recorded fEPSPs for 80 minutes. BDNF (25 ng / ml) and (LM-22A4: 5 μM) were perfused in the last 5 minutes of the baseline and in the first 5 minutes post-LTP. Signals were acquired at 100 kHz, filtered at 3 kHz (Bessel, 8-pole) with pClamp 10.2 software (Molecular Devices, USA).

### Data Analysis and Statistics

For quantitative analysis of cells positive for c-fos, two consecutive sections containing the hippocampus were taken from each animal, blindly. In each section, the number of c-Fos-positive cells was counted bilaterally using an automated cell counting procedure by ImageJ software (National Institute of Health, Bethesda, MA). The image was converted to grayscale using the thresholding feature, to highlight and standard the shape parameters of the immunoreactive particles so that the program determined what to consider a cell. Threshold settings and light intensity were kept constant across all sections and during photo acquisition. The thresholded Fos-stained cells for the area were averaged and are presented as Fos-positive nuclei/section for each brain area.

AMPA and NMDA EPSCs slope conductances were determined as the slope of linear functions of the most linear parts of the IV relationships. mIPSCs were analyzed using Mini Analysis software (Synaptosoft 6.0.3, Fort Lee, NJ, USA) and EPSCs with custom-written routines in IgorPro (Wavemetrics, Portland, OR, USA) and Matlab (MathWorks, Natick, MA, USA). The peaks of the EPSCs were used to build IV relationships to calculate the reversal potential. We measured rise times from baseline to peak and decay times from peak to baseline. mIPSCs were recorded for 10 minutes at −70 mV after the application of TTX to block action potentials and were selected manually, and only currents with good signal to noise ratio which could be easily identified as mIPSCs were chosen. Histograms for inter-event (IEI) intervals, amplitudes and rise times were built with the same fixed bins for different groups of cells and. Analysis of decay kinetics for inhibitory currents was performed by Mini Analysis group analysis with individual currents fitted with double exponential functions. Fast and slow time constants were presented as the average and compared between groups.

Field potentials were analyzed with Clampfit 10.2. Traces were low-pass filtered offline (500 Hz), and the slopes of the fEPSPs were fitted with a standard linear function. Data from each experiment were normalized relative to its baseline. LTP was quantified as the average of the fEPSP slopes in the last 20 minutes from 80 minutes post-induction recording period.

All the results are presented as means ± SE, and statistical significance was determined using unpaired Student’s t-test and one-way ANOVA as required, cumulative frequency distributions were tested for significance with Kolmogorov-Smirnov (KS) test. We use a significance level of 5% (P≤0.05).

## Results

### High-intensity sound exposure increases c-fos expression in hippocampal neurons

Because high-intensity sound affects the hippocampus in several ways (de Deus at al., 2017; Cunha et al., 2019), we tested if our protocol of high-intensity sound stimulation was able to activate hippocampal cells through of c-FOS expression. c-FOS is an immediate early gene (IEG) which expression is driven by calcium entry by NMDA receptor activation and voltage-dependent calcium channels, during intense action potential firing, and it is used as a marker of neuronal activation (Dragunow and Faull, 1989; Hudson et al., 2018). Immunostaining for c-FOS in stimulated rats showed a high level of c-FOS expression in CA1 neurons in the pyramidal layer (192.0 ± 16.71 cells/section; n=3 animals) compared to sham-stimulated (142.0 ± 3.041 cells/section; n=3 animals; P = 0.04, Figure 1A). We also observed that pyramidal layer CA3 neurons presented higher c-FOS expression in stimulated (154.8 ± 12.74 cells/section; n=3 animals) when compared sham-stimulated rats (88.50 ± 5.074 cells/section; n=3 animals; P = 0.008, Figure 1B). In contrast, c-FOS expression was similar in neurons of the dentate gyrus, in sham and stimulated rats (P ≥ 0.05, Figure 1C). These findings indicate that high-intensity sound drives robust excitatory synaptic activity, which is able to induce c-FOS expression in hippocampal neurons.

**Figure 1.**
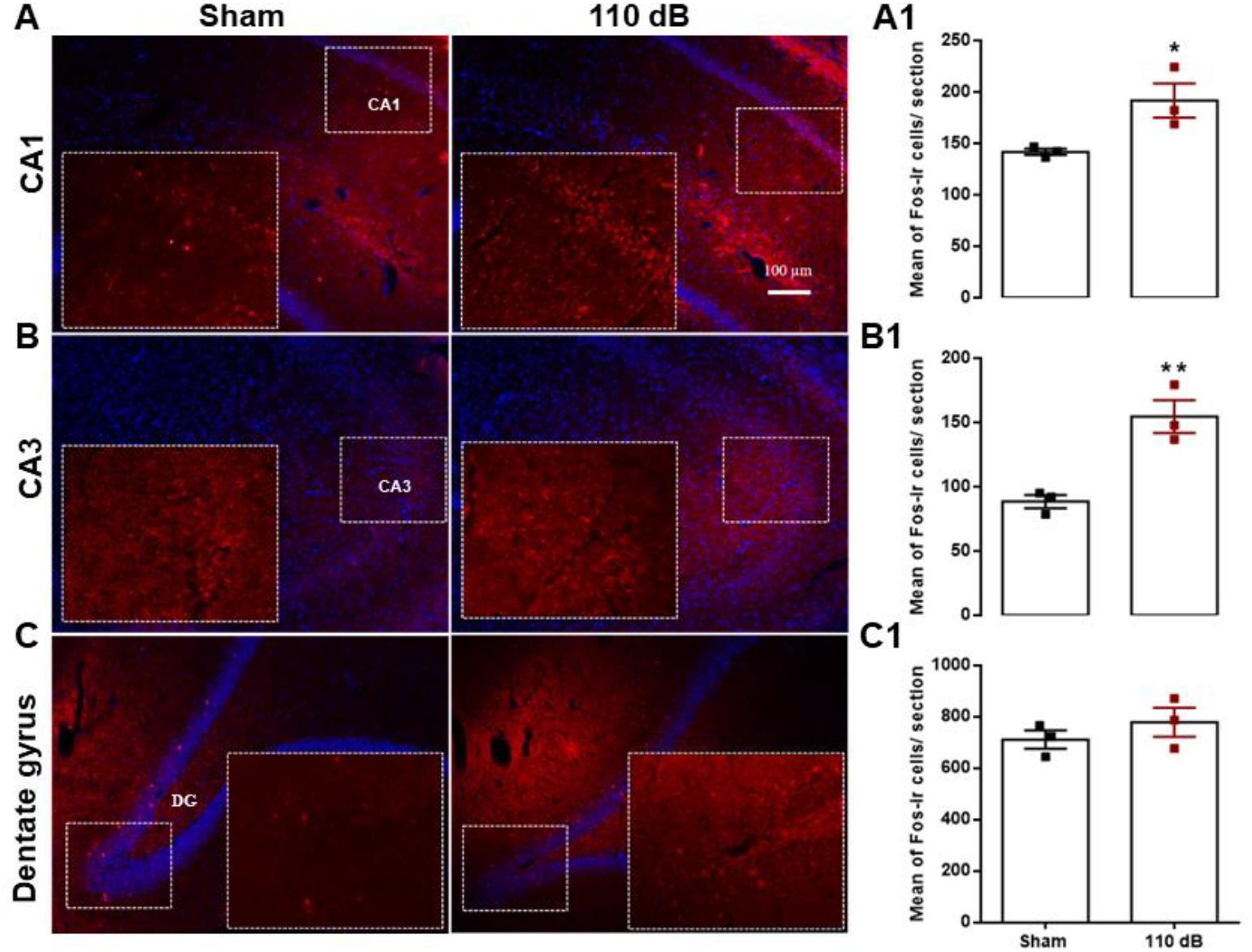
Photomicrographs illustrating c-FOS-positive nuclei labeling (red) and cell nuclei stained with DAPI (blue) in sections of CA1 (A), CA3 (B) and dentate gyrus (C). Left panel, representative images from an animal exposed to sham stimulation and an animal exposed to 110 dB sound stimulation (10X). Inset: squared area at higher magnification (20X). Right panel, the number of c-FOS-positive nuclei staining/section for CA1 (A1), CA3 (B1) and dentate gyrus (C1). *P < 0.05.

### High-intensity sound exposure does not affect glutamatergic neurotransmission in the SchafferCA1 synapses

LTP is dependent on the activation of NMDA receptors in the synapses of the Schaffer-CA1 pathway with pyramidal neurons of the hippocampus, (Collingridge and Bliss, 1987; Tang et al., 1999 and Tsien et al., 1996). The high-intensity sound induced inhibition of LTP previously shown by our group (de Deus, et al, 2017) could be due to an inhibition of NMDA-mediated currents or to a decreased AMPA/Kainate (AMPA/KA) receptor activation. Therefore, we investigated if acute high-intensity sound stimulus alters glutamatergic synaptic neurotransmission. In these experiments, we recorded AMPA/KA (DNQX-sensitive) and NMDA (DNQX-resistant) receptor-mediated EPSCs from Schaffer-CA1 synapses from sham and stimulated animals (Figure 2A). Both IV relationships of the AMPA/KA and NMDA receptor mediated EPSCs were similar between the sham and stimulated groups (Figure 2B and 2C). The calculated slope conductance of the currents was not significant different between sham and stimulated groups (AMPA/KA, sham: 8.4 ± 0.6 nS; stimulated: 8.4 ± 0.7 nS, p = 0.9; NMDA, sham: 3.8 ± 0.3 nS; stimulated: 4.0 ± 0.4 nS, p = 0.6). The amplitudes of AMPA/KA receptor-mediated EPSCs at −80 mV in sham (−616.4 ± 101.7 pA; n = 9) and stimulated animals (−657.2 ± 108.2 pA; n = 16) were not see significantly different (p = 0.80;Figure 2D). Also NMDA-mediated EPSCs at +70 mV, were not significantly different between sham (310.2 ± 56.59 pA; n = 9) and stimulated groups (370.7 ± 71.43 pA; n = 11; p = 0.52; Figure 2E). Our data show that a sound stimulus of 110 dB of 1-minute duration does not alter fast hippocampal glutamatergic neurotransmission, thus LTP inhibition is not caused by a change in the glutamatergic receptor-mediated currents in pyramidal neurons of the CA1 region.

**Figure 2.**
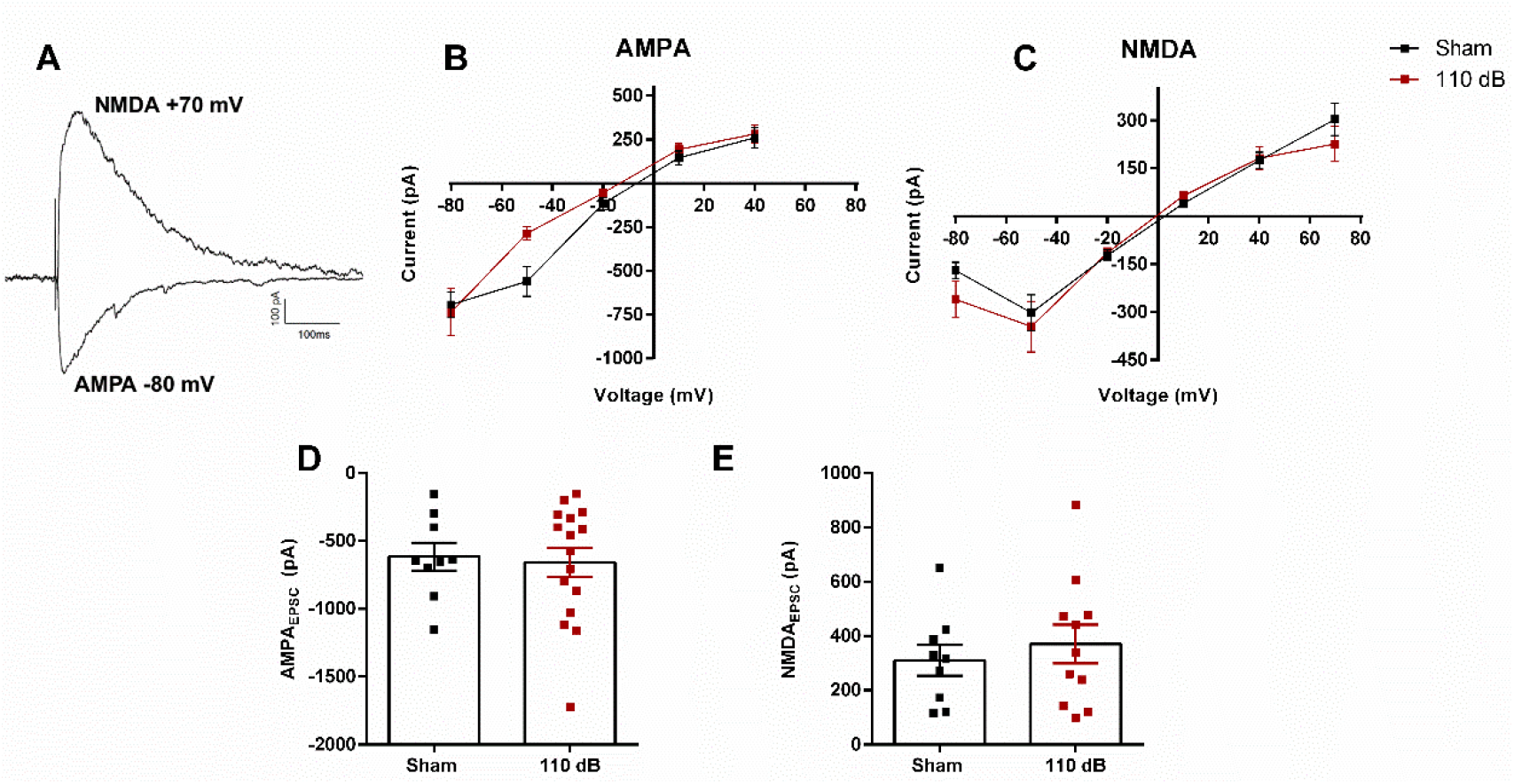
Glutamatergic neurotransmission evoked by AMPA/KA and NMDA receptors. (A) Representative traces of AMPA/KA currents (DNQX-sensitive) at −80 mV and NMDA currents (DNQX-resistant) at +70 mV, respectively. AMPA/KA (B) and NMDA (C) peak currents at different voltages. Mean peak currents of AMPA/KA (D) at −80 mV and NMDA at +70 mV (E).

### High-intensity sound exposure does not substantially affect GABAergic transmission in CA1 pyramidal neurons

Previous results from our group showed that a long-term protocol of high-intensity sound stimulation which inhibits LTP in the Schaffer-CA1 synapse (2 episodes a day, for 10 days; Cunha et al., 2015) potentiates GABAergic transmission on pyramidal CA1 hippocampal neurons (Cunha et al., 2019) by increasing the amplitude of the GABAergic currents. We then tested if a single episode of high-intensity sound could potentiate the GABAergic neurotransmission in the CA1 pyramidal neurons.

However, different to what observed with the 10-days protocol we did not observe changes in the GABAergic transmission (Figure 3A) in the hippocampus of animals subjected to one minute of high-intensity sound. Moreover, we did not find differences in the amplitude of the mIPSCs after the application of TTX: (sham stimulated: 75.77 ± 6.1 pA, n = 8; stimulated: 69.63 ± 3.52 pA, n = 8; P = 0.39, Figure 3B). We found a smaller, but no significant reduction in the frequency of the mIPSCs in neurons from stimulated animals (sham: 2.05± 0.48 Hz, n = 8, stimulated: 1.15 ± 0.16 Hz, n = 9, P = 0.08, Figure 3E). However, we found that mIPSCs from stimulated animals have faster decay times than the mIPSCs from control animals (fast decay time: sham: 2.9 ± 0.12 ms; n = 8; stimulated: 2.4 ± 0.18 ms; n = 9; P=0.04, Figure 3H; slow decay time: sham: 25.04 ± 1.22 ms, n = 8 and stimulated: 20.42 ± 0.90 ms, n=9; P = 0.003, Figure 3I). The proportion of the two components (fast/slow) was similar in both groups (sham: 53%; stimulated: 48%, Figure 3J). We conclude that, contrary to the observed after the long-term stimulation with high-intensity sound, the GABAergic inhibitory neurotransmission is not potentiated by a short stimulus with high-intensity sound, but a similar change in current kinetics was observed.

**Figure 3.**
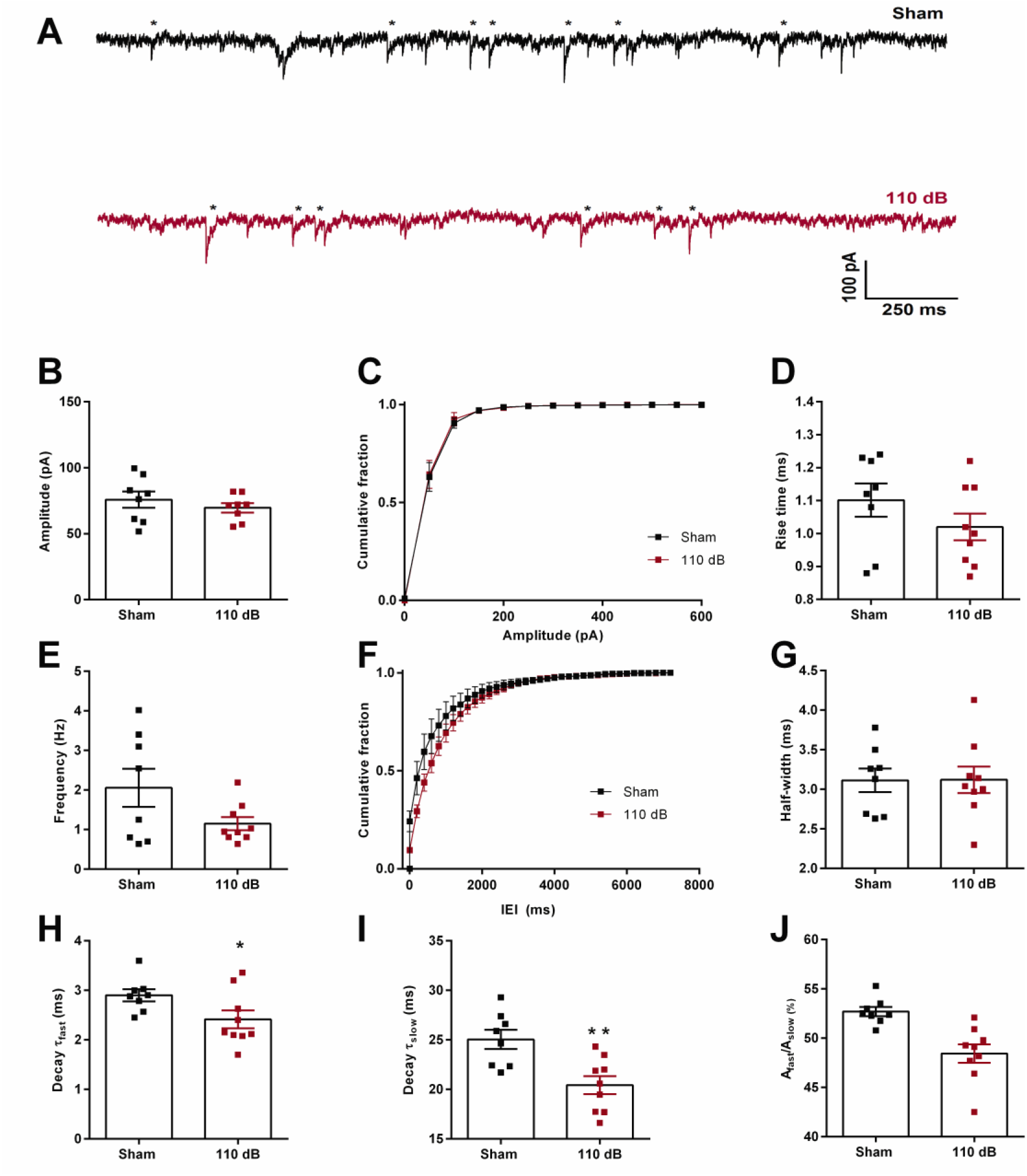
Miniature GABAergic currents. A) Representative traces of inhibitory currents from sham-stimulated rats (black trace) and rats submitted to a sound stimulus of 110 dB (red trace). B) Mean amplitude of events detected and C) cumulative fraction of amplitudes per group. D) Mean rise time of mIPSC. E) Mean frequency and F) Cumulative fraction of IEI of mIPSC. G) Mean half-widths. (H) Mean fast and (I) slow decay time constants. J) The ratio between A_fast_ and A_slow_ shown as a percentage. *P < 0.05 **P < 0.01.

**Figure 4.**
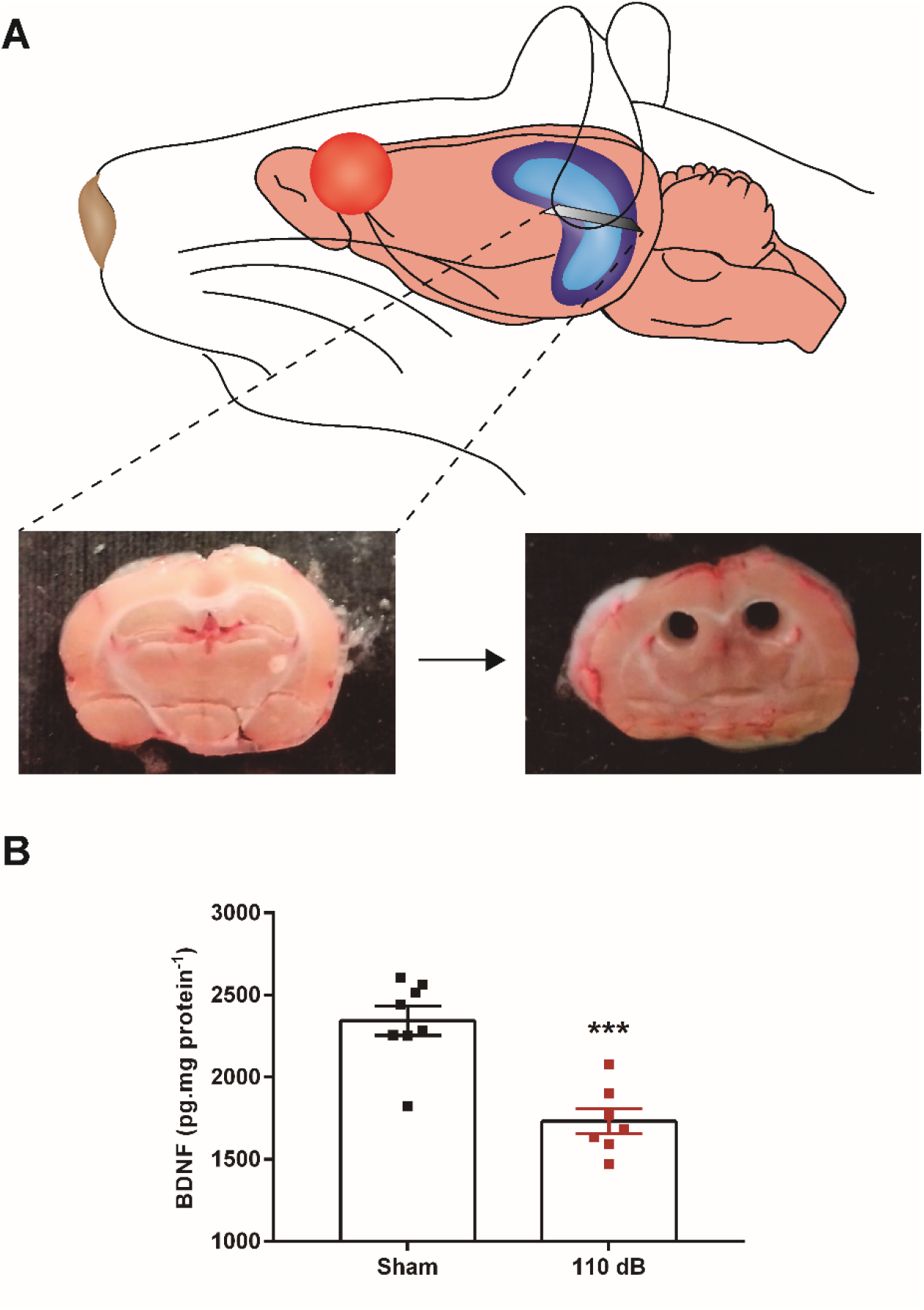
A) Schematic drawing of a transversal slice of hippocampus. B) Representative photo of bilateral dorsal hippocampus before and after the punch (1200 μm). C) BDNF levels in the dorsal hippocampus from sham-stimulated and rats submitted to a sound stimulus of 110 dB. ***P < 0.005.

### High-intensity sound exposure decreases BDNF levels in the dorsal hippocampus, and the inhibition of LTP is reverted by application of BDNF or its receptor agonist

Given that we observed no changes in the glutamatergic and inhibitory currents, we tested if another factor could account for the reduction in hippocampal LTP in these animals. Because of the strong effect of BDNF in facilitating hippocampal LTP, we hypothesized that high-intensity sound exposure could affect BDNF production or secretion in the hippocampus. Indeed, we found that BDNF levels in dorsal hippocampus are significantly decreased in stimulated (1732 ± 76.87 pg.mg protein^−1^; n=7) in comparison with sham rats (2344 ± 89.06 pg.mg protein^−1^, n=8; P=0.0002) (Figure 4). Thus, our results show that high-intensity sound decreases secretion or production of BDNF in the hippocampus.

We then decided to test if perfusion of BDNF to the hippocampal slices of rats exposed to high-intensity sound could revert the inhibition of LTP, similar to observed in the hippocampi of animal submitted to chronic intermittent hypoxia (Xie et al., 2010). First we reproduced our original observation (de Deus et al., 2017) that exposure to 110 B sound for one minute inhibits the LTP in the Schaffer-CA1 synapse (Figure 5A,B,C; sham: 1.41 ± 0.11; n = 4; sound exposed: 1.04 ± 0.06; n = 5; P = 0.01). Perfusion of BDNF (25 ng/ml) previously to the induction of LTP in the Schaffer-CA1 synapse in slices from sham animals did not change the magnitude of LTP (1.43 ± 0.07; n = 7. P = 0.90. Figure 5 D,E) showing that supplementation of BDNF to the hippocampus of animals not subjected to high-intensity sound exposure does not. However, when we perfused the slices of rats submitted to one minute of 110 dB sound with BDNF, we observed a recovery of LTP (1.55 ± 0.12; n = 6; P = 0.01) when compared to sound exposed group, to the same level of the sham group (P=0.47, Figure 5F). Additionally, we tested if the Trk-B agonist LM22A4 could restore hippocampal LTP in slices of sound-exposed rats. Indeed, perfusion of LM22A4 (5 μM) rescued hippocampal LTP from slices from sound exposed rats (1.62 ± 0.17; n = 5; P=0.02 compared with the sound exposed group) to the same level to the LTP observed in slices from sham rats; p = 0.38; Figure 5G,H). These results indicate that Trk-B receptor activation by BDNF reverts the LTP inhibition caused by exposure to high-intensity sound. Together with the reduced levels of BDNF in the hippocampus of these animals, these results suggest that exposition to on minute of high-intensity sound decreases BDNF levels in the hippocampal CA1 region resulting in inhibition of LTP induction.

**Figure 5.**
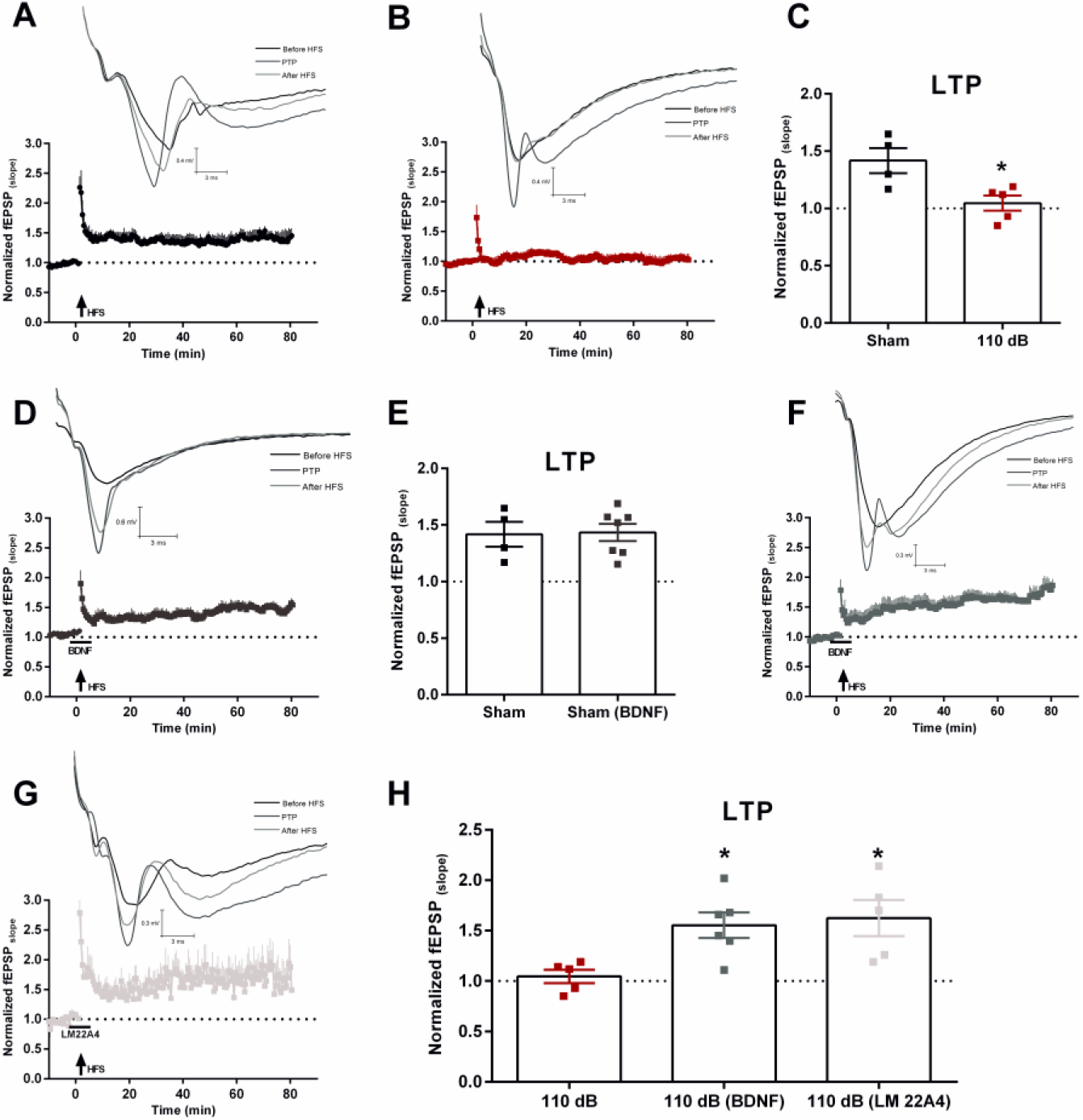
LTP from the Schaffer-CA1 synapse of sham- stimulated rats (A) and animals submitted 110 dB sound exposure (B). Normalized fEPSP slopes before and after HFS (arrow) from the Schaffer-CA1 synapse of slices of sham animals and stimulated rats. C) Bar graphs showing the summary of LTP for sham-stimulated and stimulated rats. D) Normalized fEPSP slopes before and after HFS (arrow) from the Schaffer-CA1 synapse of slices treated with BDNF (bar-10 minutes) from sham-stimulated rats. E) Bar graph showing the summary of LTP for sham-stimulated and stimulated rats. F) Normalized fEPSP slopes before and after HFS (arrow) from the Schaffer-CA1 synapses of slices treated with BDNF (bar-10 minutes) from stimulated rats. (G) Normalized fEPSP slopes before and after HFS (arrow) from the Schaffer-CA1 synapse of slices treated with LM22A4 (bar-10 minutes) from stimulated rats. Bar graph showing the summary of LTP from control slices and slices treated with BDNF and LM22A4 from stimulated rats. Representative recordings are shown above each graph. *P < 0.05.

## Discussion

Several lines of evidence show that auditory stimulation has effects beyond the auditory processing areas (Lercher et al., 2003; Stansfeld et al., 2005; Uran et al., 2010, 2012; Kraus and Canlon, 2012; Basner et al., 2014). Especially, intense auditory stimulation has been shown by us to interfere with long-term potentiation in the hippocampal Schaffer-CA1 synapse (Cunha et al., 2015; de Deus et al., 2017). Our first observation was that a 10-day long protocol of two daily episodes of 120 dB sound exposure for 1 minute each was able to inhibit the LTP in the hippocampal Schaffer-CA1 synapse of Wistar rats (Cunha et al., 2015). We chose this protocol because we were initially interested in testing if animals susceptible to audiogenic seizures from the WAR (Wistar Audiogenic Rats) strain (Doretto et al., 2003) had abnormal LTP after the protocol of audiogenic kindling (Marescaux et al., 1987; Naritoku et al., 1992; Garcia-Cairasco et al., 1996) which consisted of the exposure of the animals with the sound protocol above mentioned. To our surprise, we found that the rats from WAR strain, which consistently developed limbic seizures during the protocol of audiogenic kindling, did not have changes in LTP (Cunha et al., 2015). On the other hand, LTP in the Schaffer-CA1 synapse in the hippocampus of control Wistar rats, which did not develop audiogenic limbic seizures, was impaired (Cunha et al., 2015). We later found that a single one-minute episode of 110 dB sound was able to inhibit LTP in the Schaffer-CA1 synapse (de Deus et al., 2017), showing that exposure to high-intensity sound even for a brief period can impact hippocampal LTP. Interestingly, despite this strong effect in inhibiting LTP we did not find any deficit in spatial navigation and memory in the animals subjected to our protocols of high intensity sound exposure (Cunha et al., 2015; de Deus et al., 2017), showing that despite inhibiting hippocampal LTP, exposure to high-intensity sound does not affect basic spatial learning and memory processes.

In order to investigate the possible mechanisms of this effect on the hippocampi of Wistar rats, we performed experiments studying the synaptic transmission on the CA1 pyramidal neurons and their intrinsic electrophysiological properties (Cunha et al., 2018, 2019). The mechanisms of the inhibition of LTP by the prolonged exposure to high-intensity sound might be related, at least partially, to an increase in membrane resistance and decrease in action potential threshold, by a reduction in the expression of I_h_ in CA1 pyramidal neurons (Cunha et al., 2018) and a potentiation of GABAergic transmission (Cunha et al., 2019). However, in the animals subjected to one minute of high-intensity sound, we did not find differences in the intrinsic properties of CA1 pyramidal neurons (Cunha et al., 2018) showing that the impact of high-intensity sound in the hippocampus is dependent on the length of sound exposure.

In this work, we continued the investigation of the mechanisms of LTP inhibition after one-minute exposure to a 110 dB sound. We found like observed after prolonged exposure to one-minute episodes of 110 dB sound (Cunha et al., 2019) that the glutamatergic transmission both via AMPA/kainite and NMDA receptors are not affected by a one minute 110 dB sound exposure. However, no potentiated GABAergic transmission was observed after a single episode of 110 dB sound, differently to the potentiated GABAergic transmission after ten days of stimulation (Cunha et al., 2019). This suggests that the potentiated GABAergic transmission is a late response to high-intensity sound stimulation and probably compensates the late increased excitability of CA1 pyramidal neurons observed after the prolonged exposure to high-intensity sound (Cunha et al., 2018). Thus, this indicates that the effects of high-intensity sound are distinct in both models of high-intensity sound exposure.

The hippocampus is connected to the auditory system indirectly from the frontomedial cortex, insula, and amygdala (Kraus and Canlon, 2012), by a pathway from the cochlear nucleus to the entorhinal cortex involving the pontine reticula nucleus, pontine central gray and medial septum, with a higher threshold than the canonical pathway via inferior colliculus (Zhang et al., 2018) and by a direct connection between the auditory cortex and the CA1 region (Zhao et al., 2018). This pathway is implicated in the formation of long-term auditory and auditory-spatial memories (Squire et al., 2001; Tamura et al., 1990; Arononv et al., 2017) and fear conditioning (Zhang et al., 2018). Accordingly, we showed that c-FOS expression is increased in the pyramidal cell layer of both CA1 and CA3 regions. C-FOS expression is induced by strong increases in intracellular calcium trough NMDA receptors and voltage-dependent calcium channels, which activates MAPK pathway leading to the phosphorylation of the transcription factors CREB and Elk-1, that bind to the c-fos promoter (Chung, 2015; Hudson, 2018). The expressed FOS protein dimerizes with the protein JUN to form the AP-1 complex that regulates the expression of several genes that could be involved in the inhibitory effect of high-intensity sound on LTP. Because c-fos expression needs strong calcium influx induced by synaptic activity, we could hypothesize that high-intensity sound stimulation induces a hyperexcitation on hippocampal neurons during the exposure to sound. We found increased c-fos expression in the pyramidal layer, were the glutamatergic pyramidal neurons are located, however we cannot discard the activation of GABAergic interneurons located in the pyramidal layer or adjacent to it. Interestingly, *in vivo* hippocampal recordings showed that sound exposure evokes inhibitory currents on pyramidal neurons, accordingly to the activation of hippocampal inhibitory interneurons (Wang et al., 2017).

Based on the critical role of BDNF over hippocampal LTP (Minichielo, 2009; Edelmann et al., 2014; Lin et al., 2018), we tested the hypothesis that a lack of BDNF could be related to the LTP impairment after high-intensity sound exposure. We found that the expression of the mature form of BDNF was reduced in the hippocampus of rats exposed to high-intensity sound. This could represent a depletion of the cellular BDNF and/or increasing degradation of existing BDNF. We then demonstrated that exogenous application of BDNF was able to rescue the LTP from the Schaffer-CA1 synapses from rats exposed to high-intensity sound. Because BDNF did not increase the magnitude of LTP from the Schaffer-CA1 synapses from sham rats, the increase in the magnitude of LTP in sound exposed animals was not due to a facilitation of LTP by exogenous BDNF, but we believe it represented a replenishment of reduced BDNF levels in the hippocampus of these animals. Additionally, the agonist of the Trk-B receptor LM22A4 also rescued the LTP from sound stimulated animals, confirming that the hippocampal BDNF/ Trk-B pathway is affected by high-intensity sound exposure. However, the efficacy of LM22A4 and other small molecule agonists of Trk-B receptors has been recently questioned (Boltaev et al., 2017). Thus, we conclude that a decrease in BNDF levels in the hippocampus of rats submitted to one minute of high-intensity sound, impairs the expression of LTP in the Schaffer-CA1 synapse. Interestingly, Matt et al., (2018) found that prolonged exposure to traumatic noise (120 dB, 6 hours per day for 21 days) in anesthetized rats did not alter LTP and BDNF expression. This suggests that the anesthesia or the prolonged sound exposure protocol might change the hippocampal response to high-intensity sound. We do not know, however, if the same decrease in BDNF levels occurs in the hippocampus of animals subjected to the 10-days long sound exposure protocol.

Our results add to other reports showing the influence of the acoustic environment on hippocampal function. Both moderate and intense sound exposure affect hippocampal function in several ways. For instance prolonged (2 hours/ day for 3–6 weeks) moderate (80 dB) sound exposure, impairs spatial memory in mice and increases oxidative damage and hippocampal tau phosphorylation (Cheng et al., 2011, 2016). On the other hand, daily exposure to music at moderated intensity levels (60 dB, 6 hours per day for 21 days) enhanced learning performance and increased BDNF expression in the hippocampus (Angelucci et al., 2007). Similarly, exposure to an 80 dB sound at 10 kHz for 40 minutes in anesthetized rats, increased BDNF expression, hippocampal LTP magnitude and improved the performance in the Morris water maze (Matt et al., 2018). It appears that the effect of moderate noise is more related to the type of noise, and it seems to be contradictory depending on the reports. Exposure to intense noise, on the other hand, is often associated with deficits in hippocampal function. For instance, acute traumatic noise (106–115 dB, 30–60 minutes) alters place cell activity in the hippocampus and increases arc expression, an immediate early gene related to synaptic plasticity, in the hippocampi of rats. Prenatal exposure to loud sounds has a deleterious effect on the hippocampal LTP hippocampal-dependent learning and memory in rats (Barzegar et al., 2015). Sound deprivation has also been shown to impact hippocampal function. Zhao et al., (2018) showed that temporary conductive hearing loss in young rats reduced LTP, NMDA currents, dendritic spine density and resulted in impaired performance in the Morris water maze 30 days later when the hearing was restored.

Thus, the results presented here and in the other reports from our group, in combination with the above-mentioned evidences, show that the hippocampus is strongly influenced by the auditory environment and that the BDNF is an important mediator of these effects. The knowledge about the influence of noise exposure on non-auditory areas like the hippocampus is extremely relevant because exposure to loud noises is an increasingly common occurrence in our daily life, and several cognitive and emotional deficits are associated to prolonged and acute noise exposure in humans (Lercher et al., 2003; Stansfeld et al., 2005; Basner et al., 2014; Hofner et al., 2018; Swanson et al., 2018).

## Acknowledgments

Work supported by São Paulo State Research Foundation (FAPESP) grants 2015/22327-7, 2016/01607-4 and 2016/17681-9. We thank the technical assistance of Mr. J. Fernando Aguiar and Mr. Rubens Fernando de Melo.

